# KmerSV: a visualization and annotation tool for structural variants using Human Pangenome derived k-mers

**DOI:** 10.1101/2023.10.11.561941

**Authors:** Qingxi Meng, Hanlee P. Ji, HoJoon Lee

**Affiliations:** Division of Oncology, Department of Medicine – Stanford University School of Medicine CCSR 1115, 269 Campus Drive

## Abstract

**Summary:** KmerSV is a visualization and annotation tool for structural variants (**SVs**). It can be applied to assembly contigs or long-read sequences. Using k-mers it rapidly generates images and provides genome features of SVs. As an important feature, it utilizes the new Human Pangenome reference which provide haploid specific assemblies, addresses limitations in prior references and improves the discovery of SVs.

**Availability and implementation:** KmerSV is implemented in Python and available at github.com/sgtc-stanford/kmerSV

## 1 Introduction

Structural variants (**SVs**) are large rearrangements that include deletions, insertions, inversions, duplications among others. SVs contribute to a variety of human diseases (Beyter, et al., 2021; Feuk, et al., 2006) such as Mendelian disorders (Sanchis-Juan, et al., 2018), cancer (Feuk, et al., 2006), neurological conditions (Vialle, et al., 2022), and cardiovascular maladies (Costain, et al., 2016). Recent genomic advances such as long-read sequencing improve the identification and characterization of SVs (Gong, et al., 2021; Ho, et al., 2020; Mastrorosa, et al., 2023; Mitsuhashi and Matsumoto, 2020). With the increasing number of sequenced whole genomes, there is a great need for efficient, straightforward visualization to examine, validate and generate images of SVs (Krzywinski, et al., 2009; Mitsuhashi and Matsumoto, 2020). Tools such as the UCSC Genome Browser or the Integrative Genomics Viewer (**IGV**) provide tools for visualizing variants. However, these applications are limited when it comes to visualizing SVs.

We introduce KmerSV, a new tool for SV visualization and annotation. To mediate these functions, KmerSV uses a reference sequence deconstructed into its component k-mers, each having a length of 31 bp. These reference-derived k-mers are compared to the sequence of interest. For this application, we leveraged National Institute of Health’s recently released Human Pangenome reference (Lee, et al., 2023; Liao, et al., 2023). The program maps the Pangenome or other reference 31-mers against one or multiple target sequences which can include either contigs or sequence reads. Initially, we retrieve these k-mers via a sliding window across a segment of the reference with its coordinate information. Then, the retrieved k-mers are systematically mapped against the target (i.e., contig or read). Generally, a portion will be uniquely mapped to the target sequence. However, some k-mers will map to multiple locations as commonly seen in duplications or highly repetitive sequences. Unique 31-mers (as defined by the reference) serve as “anchor” points in the target sequence to facilitate using k-mers with multiple coordinates. This anchoring process eliminates ambiguous k-mers and improves the visualization of complex SVs such as duplications. KmerSV also incorporates gene and exon annotations within its visual display, enabling straightforward identification of the specific genomic elements associated with each SV.

KmerSV leverages the Human Pangenome reference. For its first release this new genome standard based on 94 haploid genomes sourced from 47 diverse individuals. This new reference provides haploid features, fills in gaps and has greater representation of human genetic diversity compared to the prior reference GRCh38 (Liao, et al., 2023). From our previous work, we identified the unique k-mers from the Human Pangenome (Lee, et al., 2023). Then, KmerSV visualizes the SV in the context of the component haploid assemblies making up the Human Pangenome.

### 2 KmerSV

To provide SV images and annotation, the program has several steps: 1) assembling a table that lists the sequences of 31-mers from a given reference; 2) determining their respective coordinates in from both the reference and target sequence; 3) graphically representing these 31-mer positions using a scatter plot; 4) overlaying relevant genomic annotations to enhance the resulting visualization (**Figure 1**). KmerSV uses two sequences: a reference sequence such as the Human Pangenome and a target sequence. The image involves pairing the reference sequence with several target sequences. For genome feature annotation of the SV, users can optionally input a custom gene annotation BED (Browser Extensible Data) file pertinent to the reference sequence. If a BED file is not supplied, KmerSV defaults to an annotation file derived from the Matched Annotation from NCBI and EMBL-EBI (**MANE**) (Morales, et al., 2022).

**Figure 1.**
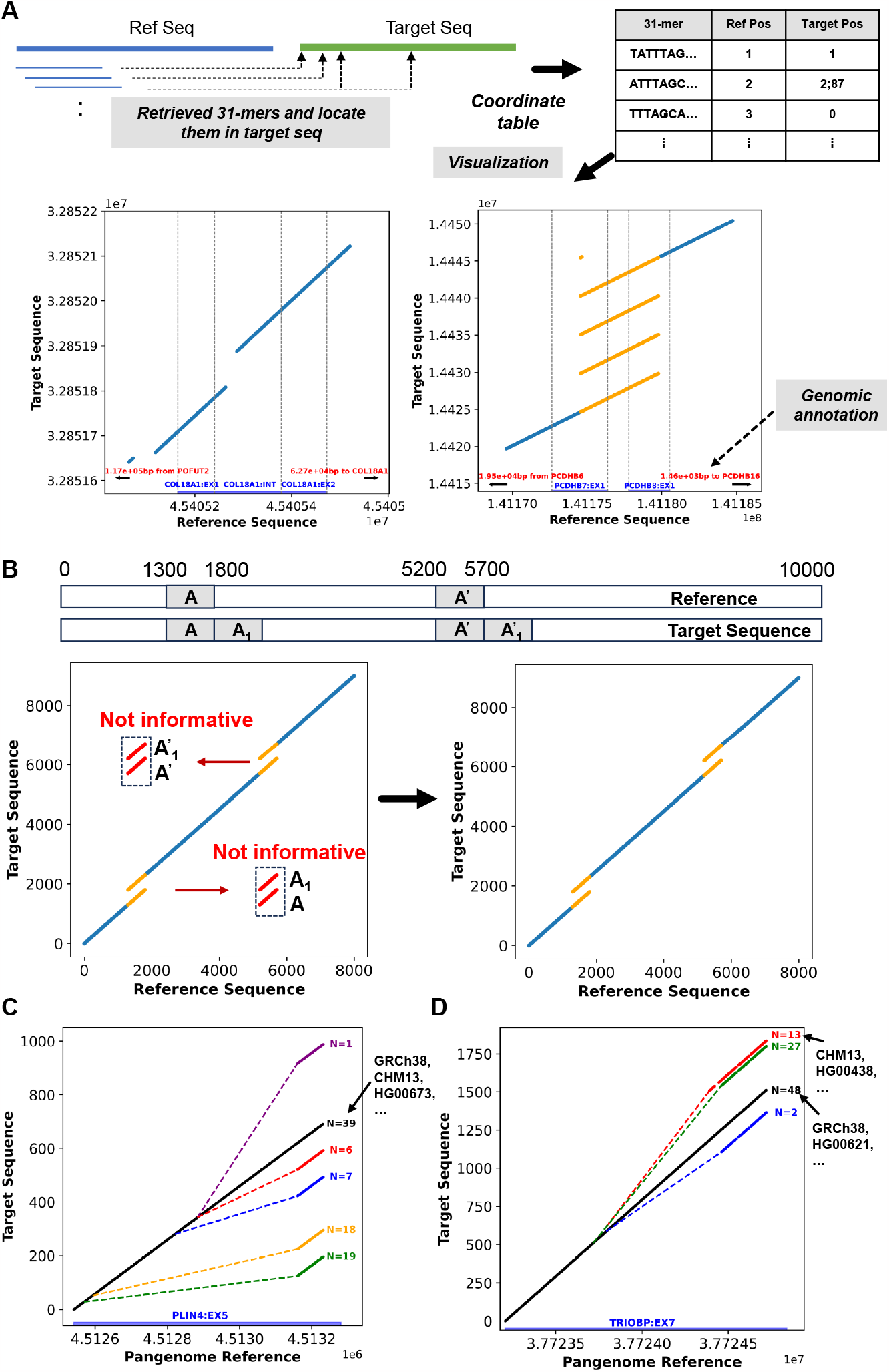
Overview and examples of the KmerSV framework. (A) Overview of KmerSV analysis. In the plots, unique 31-mers are shown in blue, while non-unique 31-mers are shown in orange. (B) Process of resolving ambiguous, non-unique 31-mers: A and A’ represent identical sequences, as do A1’ and A1’. The unique 31-mers are highlighted in blue, relevant non-unique 31-mers in orange, and irrelevant ones in red. (C) & (D) Visualization of SVs in the pangenome reference graph. In Figure C and D, the black line represents the reference sequence and all of the Human Pangenome haploid references that align perfectly with it. Each distinct colored line depicts a structural deviation from the selected reference sequence. For example, in Figure 1C, the black line corresponds to GRCh38 and specific Human Pangenome haploid references like HG00673 that align perfectly. The purple line indicates an insertion approximately 350bp from the reference sequence’s beginning, observed in just one of the 90 haploids. The orange line indicates a deletion around 50bp from the reference sequence’s beginning, observed in 18 of the 90 haploids. The red, blue, and green lines mark three other distinct deletions present in certain haploids.

### 2.1 Generating a k-mer coordinate table

KmerSV retrieves 31-mers from the reference sequence (i.e., the Human Pangenome) with its coordinate position. We also provide the option to use the GRCh38 or other references. Subsequently, each k-mer and its reverse complement are mapped onto the target sequence. Notably, KmerSV employs regular expressions to determine the position of exact matches for each k-mer. The output is a tabular compilation of the informative 31-mers present in both the reference and target sequence and with their coordinate position and strand (**Fig. 1A**).

Typically, a one-to-one match is anticipated for unique 31-mers found in both the reference and the target sequence. The coordinates of these matches are provided in the table for both the reference and target sequences. If a 31-mer from the reference lacks a match in the target sequence because of deletions or single nucleotide polymorphisms (**SNPs**), its entry value in the table is 0. When a 31-mer maps to several locations on the target sequence, these coordinates are captured in the table and delineated by semicolons (;).

### 2.2 Visualization with annotations

KmerSV utilizes the k-mer coordinate table for visualizing SVs. The x-axis represents the coordinates of the 31-mers in the reference sequence, whereas the y-axis provides their positions in the assembly contig or sequence read. A diagonal line should manifest when there are no SVs in the target sequence. Any deviations from the diagonal suggest the presence of SVs. For example, an insertion leads to a vertical gap on the Y axis of the plot. On the other hand, a deletion leads to a horizontal gap on the X axis of the plot. We present two SV visualization examples in **Figure 1A**. The left plot shows a deletion (at coordinate bp 45,405,100) and an insertion (at coordinate bp 45,405,260). The right plot clearly reveals a segmental duplication with four copies (in the vicinity of coordinate bp 141,174,553). Detailed sequence information can be found in the **Supplementary Material**.

KmerSV integrates gene, exon, and intron annotations into its visualization (**Figure 1A**). It utilizes Python package pandas to identify overlapping regions between the provided (or default) BED file and identified SVs. Relevant information from overlapping regions, whether gene, exon, or intron, is merged into the plot, enhancing the visual representation. For intergenic regions of interest, the tool also provides distances to adjacent genes, offering an informative genomic context for each SV.

### 3 Filtering ambiguous 31-mers

There are examples of complex SVs when mapping k-mers between a reference and target sequence generate a convoluted graphic plot that does not reveal the structure. To provide a clear representation of these type of SVs, we eliminate non-informative k-mers with a two-step method. First, we display only unique 31-mers as determined from the reference. Next, we include non-unique k-mers that are adjacent to the unique ones. Figure 1B shows the unique 31-mers as blue dots, adjacent non-unique 31-mers as orange dots, and distal, non-informative 31-mers as red dots. Then, KmerSV incorporates a distance threshold and multi-location filtering to eliminate ambiguous 31-mers. This procedure includes only those non-unique k-mers within 1 kb of the unique k-mers. This is the default setting. Second, those k-mers which are highly repetitive and with 100 or more multiple coordinate positions are eliminated. This filtering approach allows KmerSV to consider SVs even in the presence of k-mers with SNPs or ones that occur in repetitive sequences. This two-step approach, coupled with a preset threshold, is robust and computationally efficient compared to other methods as detailed in the **Supplementary Materials**.

### 4 Structural Variants Within the Framework of the Human Pangenome Reference

The Human Pangenome is a compilation of multiple haploid assemblies from different individuals. As a result, there is variation in the SV structure among the various haploid assemblies. **Figure 1C and D** showcase how the KmerSV tool in delineating these differences when applied to the Human Pangenome graph. We downloaded the most recent pangenome graph for chr19 from the human-pangenomics S3 bucket (https://s3-us-west-2.amazonaws.com/human-pangenomics/index.html) (Liao, et al., 2023). This graph data encompasses a total of 90 haplotyped assemblies. Subsequently, we employed odgi (v.0.6.2) (Guarracino, et al., 2022) to extract all paths within the region GRCh38.chr19: 4512541 - 4513161 from the graph, using the example commands provided below:

odgi extract -i chr19.full.og -o chr19.region.og -b chr19.region.bed -E –P

odgi paths -i chr19.region.og -f > chr19.region.fasta

Next, we used the KmerSV tool on specific haploid sequences to transpose genome annotation that includes features such as exons. The resulting plot juxtaposes the reference sequence with the target sequence’s variations. The x-axis delineates the coordinates and annotation as per the GRCh38 reference, while the y-axis denotes the starting position of each extracted haploid assembly, with ‘0’ marking the initial position for every sample. This visualization precisely identified the specific types of SVs present across various genomes.

As illustrated in **Figure 1C**, the solid black line represents the various Human Pangenome haploid assemblies that align to the GRCh38 reference. Each distinct colored line depicts a structural change compared to the GRCh38 sequence. KmerSV represents these differences with dashed lines, connecting the SV’s start to its endpoint. Additionally, the number of haploids sharing a specific SV is annotated adjacent to the related colored line. The list of haploids, along with their corresponding colors, is provided in the output text file (see Supplementary Material). For example, the purple line indicates an insertion approximately 350bp observed in just one of the 90 haploids. The plot also showed four unique deletions distributed across multiple haploids. Notably, KmerSV also provides a gene annotation on the x-axis, emphasizing that the detected SVs are positioned within the fifth exon of the *PLIN4* gene.

**Figure 1D** showcases another instance, highlighting all samples from the region chr22: 37723714-37724734 extracted from the Pangenome graph and compared to the GRCh38 sequence. The visualization reveals an insertion and two unique deletions, all situated within the seventh exon of the *TRIOBP* gene. The GRCh38 and CHM13 sequences are different in this region, characterized specifically by CHM13 having an insertion of CHM13 compared to GRCh38. Similarly, the output text file provides the list of haploids and their associated colors (**Supplementary Material**).

## 4 Discussion and Conclusion

We present KmerSV, a computational tool designed for the rapid visualization and annotation of SVs. By employing filtering techniques, KmerSV provides an accessible and clear SV representation in given sequences. This tool offers an intuitive visualization of haploid genome assemblies in a specified region within the Human Pangenome graph. KmerSV is very rapid in generating outputs: it processes and annotates a 10 kb sequence pair within 5 seconds. Further insights into its speed and scalability are provided in the **Supplementary Material**. In the future, we will integrate KmerSV with other alignment tools. This would facilitate the automatic identification of sequences or reads potentially containing SVs, such as those indicated by the CIGAR string, as primary inputs.

## Funding

All authors were supported by National Institutes of Health grant [U01HG01096]. HPJ received additional support from the Clayville Foundation.

## Notes

https://github.com/sgtc-stanford/kmerSV

### Competing Interest Statement

The authors have declared no competing interest.

