## Supplementary Materials for "KmerSV: a visualization and annotation tool for structural variants using Human Pangenome derived k-mers"

### Supplementary material for KmerSV

#### Details for the visualization plots in Figure 1A in the main paper

The left plot employs GRCh38 chr21:45405064-45405552 as the reference sequence, with HG002 maternal JAHKSD010000059.1:32851608-32852153 serving as the target sequence. The right plot utilizes GRCh38 chr5:141169588-141184761 as its reference sequence, paired with HG02080 maternal chr5 141174588-141179791 as the corresponding target sequence.

#### Details about filtering of ambiguous 31-mers

Ambiguous 31-mers are sequences that may be either erroneously aligned or mapped to multiple sites within the target sequence. Such ambiguities can arise due to factors like single nucleotide polymorphisms (SNPs), which may result in the incorrect alignment of specific 31-mers from the reference genome to the target. Additionally, the existence of repetitive 31-mers in both reference and target sequences can further muddle the visualization process due to their multiple locations. These problematic 31-mers often obscure the clear visualization of structural variations (SVs). Therefore, our aim is to filter out these ambiguous 31-mers while maintaining the accuracy of the remaining, unique 31-mers.

Initially, we hypothesized that low-complexity 31-mers, such as those containing 20 adenines, would predominantly appear among repetitive 31-mers. To test this, we calculated both the Shannon index and Kolmogorov complexity for each 31-mer extracted from the reference sequence. We then compared the distribution of these indices between unique 31-mers and ambiguous, non-unique ones. Contrary to our expectations, the distributions were strikingly similar, indicating that these metrics were not effective for filtering out ambiguous 31-mers. The limitations of the Shannon index—measuring only nucleotide frequency without accounting for sequence order—became evident. While Kolmogorov complexity does consider sequence order, the variable complexities of ambiguous 31-mers induced by SNPs made setting a definitive threshold difficult.

We also experimented with generating a blacklist for 31-mers that appear more than 100 times across the reference genome (GRCh38), with the intent of excluding these from the visualization. However, this strategy turned out to be ineffective in practice. It became clear that 31-mers that are globally repetitive aren't necessarily locally repetitive, undermining the utility of a global blacklist in refining our visualization.

In the end, we opted not to employ any of the previously mentioned methods in our implementation. Instead, we found that setting an appropriate distance threshold for discarding 31-mers that are far removed from their neighboring unique 31-mers provided adequate filtration of ambiguous 31-mers. While more sophisticated approaches that take into account local sequence complexity might potentially eliminate additional ambiguous 31-mers, the computational overhead of these methods would be significantly higher.

##### Details for the pangenome example in Figure 1C and D

In the main paper's Figure 1C, each sample is denoted by a line of a distinct color. Comprehensive details about each sample, along with its corresponding line color, are available in Supplementary Tables 1 and 2.

Supplementary Table 1: Detailed Information about the samples in left plot of the Figure 1C.

| Assembly ID | Contig ID | Haplotype | Start Pos | End Pos | Color in Figure 1D |
|---|---|---|---|---|---|
| GRCh38 | chr19 | - | 4512541 | 4513263 | Black |
| CHM13 | chr19 | - | 4496224 | 4496946 | Black |
| HG00438 | JAHBCB010000083.1 | Paternal | 1885724 | 1886446 | Black |
| HG00438 | JAHBCA010000108.1 | Maternal | 1882886 | 1883608 | Black |
| HG00621 | JAHBCD010000127.1 | Paternal | 2097633 | 2098355 | Black |
| HG00621 | JAHBCC010000085.1 | Maternal | 3480601 | 3481323 | Black |
| HG00673 | JAHBBZ010000102.1 | Paternal | 2535453 | 2536175 | Black |
| HG00673 | JAHBBY010000058.1 | Maternal | 2539139 | 2539861 | Black |
| HG00733 | JAHEPQ010000028.1 | Paternal | 20245062 | 20245784 | Black |
| HG00733 | JAHEPP010000036.1 | Maternal | 3482029 | 3482652 | Red |
| HG00735 | JAHBCH010000086.1 | Paternal | 21122573 | 21122800 | Green |
| HG00735 | JAHBCG010000124.1 | Maternal | 3361690 | 3362016 | Orange |
| HG00741 | JAHALY010000106.1 | Paternal | 1183181 | 1183804 | Red |
| HG00741 | JAHALX010000059.1 | Maternal | 2716594 | 2717316 | Black |
| HG01071 | JAHBCF010000220.1 | Paternal | 195083 | 195805 | Black |
| HG01071 | JAHBCE010000080.1 | Maternal | 2536728 | 2536955 | Green |
| HG01106 | JAHAMC010000044.1 | Paternal | 3491013 | 3491735 | Black |
| HG01106 | JAHAMB010000014.1 | Maternal | 35139636 | 35140358 | Black |
| HG01109 | JAHEPA010000086.1 | Paternal | 2532809 | 2533531 | Black |
| HG01109 | JAHEOZ010000187.1 | Maternal | 2541231 | 2541557 | Orange |
| HG01123 | JAGYYZ010000080.1 | Paternal | 3482141 | 3482863 | Black |
| HG01123 | JAGYYY010000010.1 | Maternal | 2103255 | 2103977 | Black |
| HG01175 | JAHAMA010000167.1 | Paternal | 2146565 | 2146792 | Green |
| HG01175 | JAHALZ010000117.1 | Maternal | 2144118 | 2144840 | Black |
| HG01243 | JAHEOY010000101.1 | Paternal | 2532959 | 2533681 | Black |
| HG01243 | JAHEOX010000170.1 | Maternal | 1658280 | 1658606 | Orange |
| HG01258 | JAGYYV010000123.1 | Paternal | 10543422 | 10544144 | Black |
| HG01258 | JAGYYU010000096.1 | Maternal | 2096432 | 2097154 | Black |
| HG01358 | JAGYZB010000117.1 | Paternal | 2143022 | 2143744 | Black |
| HG01358 | JAGYZA010000093.1 | Maternal | 14016939 | 14017661 | Black |
| HG01361 | JAGYYX010000050.1 | Paternal | 1184825 | 1185448 | Red |
| HG01361 | JAGYYW010000029.1 | Maternal | 1189984 | 1190706 | Black |
| HG01891 | JAGYVO010000075.1 | Paternal | 4323686 | 4323913 | Green |

| | | | | | |
|---|---|---|---|---|---|
| HG01891 | JAGYVN010000038.1 | Maternal | 20785165 | 20785491 | Orange |
| HG01928 | JAGYVQ010000044.1 | Paternal | 2532441 | 2533163 | Black |
| HG01928 | JAGYVP010000101.1 | Maternal | 1880042 | 1880764 | Black |
| HG01952 | JAHAME010000096.1 | Paternal | 9797458 | 9798180 | Black |
| HG01952 | JAHAMD010000107.1 | Maternal | 3461632 | 3462255 | Red |
| HG01978 | JAGYVS010000079.1 | Paternal | 2535626 | 2536348 | Black |
| HG01978 | JAGYVR010000097.1 | Maternal | 1880353 | 1881075 | Black |
| HG02055 | JAHEPK010000066.1 | Paternal | 2573270 | 2573497 | Green |
| HG02055 | JAHEPJ010000093.1 | Maternal | 2536997 | 2537224 | Green |
| HG02080 | JAHEOW010000231.1 | Paternal | 1880606 | 1881328 | Black |
| HG02080 | JAHEOV010000135.1 | Maternal | 1880124 | 1880846 | Black |
| HG02109 | JAHEPG010000011.1 | Paternal | 2539018 | 2539344 | Orange |
| HG02109 | JAHEPF010000211.1 | Maternal | 2542498 | 2543121 | Red |
| HG02145 | JAHKSG010000108.1 | Paternal | 2531665 | 2531991 | Orange |
| HG02145 | JAHKSF010000140.1 | Maternal | 4505836 | 4506558 | Black |
| HG02148 | JAHAMG010000109.1 | Paternal | 3031151 | 3031873 | Black |
| HG02148 | JAHAMF010000029.1 | Maternal | 3420787 | 3421509 | Black |
| HG02257 | JAGYVI010000050.1 | Paternal | 1879687 | 1880013 | Orange |
| HG02257 | JAGYVH010000108.1 | Maternal | 2101154 | 2101381 | Green |
| HG02486 | JAGYVM010000025.1 | Paternal | 21785583 | 21785810 | Green |
| HG02486 | JAGYVL010000047.1 | Maternal | 2139812 | 2140138 | Orange |
| HG02559 | JAGYVK010000002.1 | Paternal | 50303120 | 50303644 | Blue |
| HG02559 | JAGYVJ010000110.1 | Maternal | 1183377 | 1183604 | Green |
| HG02572 | JAHAOW010000452.1 | Paternal | 582156 | 582680 | Blue |
| HG02572 | JAHAOV010000209.1 | Maternal | 2103361 | 2103588 | Green |
| HG02622 | JAHAOO010000021.1 | Paternal | 3477772 | 3478494 | Black |
| HG02622 | JAHAON010000085.1 | Maternal | 3363119 | 3363643 | Blue |
| HG02630 | JAHAOQ010000120.1 | Paternal | 3484501 | 3484827 | Orange |
| HG02630 | JAHAOP010000098.1 | Maternal | 4510978 | 4511205 | Green |
| HG02717 | JAHAOS010000082.1 | Paternal | 2141717 | 2142043 | Orange |
| HG02717 | JAHAOR010000110.1 | Maternal | 2131900 | 2132226 | Orange |
| HG02723 | JAHEOU010000153.1 | Paternal | 2143627 | 2143854 | Green |
| HG02723 | JAHEOT010000239.1 | Maternal | 2144310 | 2144537 | Green |
| HG02818 | JAHEOS010000049.1 | Paternal | 2146466 | 2146693 | Green |
| HG02818 | JAHEOR010000031.1 | Maternal | 2532086 | 2532313 | Green |
| HG02886 | JAHAOU010000021.1 | Paternal | 36598725 | 36599744 | Black |
| HG02886 | JAHAOT010000083.1 | Maternal | 2144166 | 2144888 | Purple |
| HG03098 | JAHEPM010000150.1 | Paternal | 2532251 | 2532775 | Blue |
| HG03098 | JAHEPL010000061.1 | Maternal | 1884205 | 1884828 | Red |
| HG03453 | JAGYVW010000134.1 | Paternal | 2498281 | 2498508 | Green |
| HG03453 | JAGYVV010000123.1 | Maternal | 4496413 | 4496739 | Orange |

| | | | | | |
|---|---|---|---|---|---|
| HG03486 | JAHEOQ010000045.1 | Paternal | 2544121 | 2544447 | Orange |
| HG03486 | JAHEOP010000140.1 | Maternal | 2535875 | 2536201 | Orange |
| HG03492 | JAHEPI010000200.1 | Paternal | 2141596 | 2142318 | Black |
| HG03492 | JAHEPH010000234.1 | Maternal | 2140308 | 2141030 | Black |
| HG03516 | JAGYYT010000054.1 | Paternal | 1145574 | 1145900 | Orange |
| HG03516 | JAGYYS010000062.1 | Maternal | 2306529 | 2306855 | Orange |
| HG03540 | JAGYVY010000103.1 | Paternal | 4522006 | 4522233 | Green |
| HG03540 | JAGYVX010000155.1 | Maternal | 3474409 | 3474636 | Green |
| HG03579 | JAGYVU010000032.1 | Paternal | 2531128 | 2531454 | Orange |
| HG03579 | JAGYVT010000202.1 | Maternal | 3479999 | 3480325 | Orange |
| NA18906 | JAHEOO010000085.1 | Paternal | 2139016 | 2139540 | Blue |
| NA18906 | JAHEON010000045.1 | Maternal | 2552850 | 2553374 | Blue |
| NA20129 | JAHEPE010000213.1 | Paternal | 2535650 | 2536372 | Black |
| NA20129 | JAHEPD010000068.1 | Maternal | 2526303 | 2526827 | Blue |
| NA21309 | JAHEPC010000262.1 | Paternal | 3665319 | 3665546 | Green |
| NA21309 | JAHEPB010000151.1 | Maternal | 2524217 | 2524939 | Black |

Supplementary Table 2: Detailed Information about the samples in the Figure 1D.

| Assembly ID | Contig ID | Haplotype | Start Pos | End Pos | Color in Figure 1D |
|---|---|---|---|---|---|
| GRCh38 | chr22 | - | 37723207 | 37724750 | Black |
| CHM13 | chr22 | - | 38184422 | 38186259 | Red |
| HG00438 | JAHBCB010000005.1 | Paternal | 30958648 | 30960485 | Red |
| HG00438 | JAHBCA010000050.1 | Maternal | 13123328 | 13125159 | Green |
| HG00621 | JAHBCD010000015.1 | Paternal | 386570 | 388401 | Green |
| HG00621 | JAHBCC010000017.1 | Maternal | 386320 | 387863 | Black |
| HG00673 | JAHBBZ010000029.1 | Paternal | 383958 | 385789 | Green |
| HG00673 | JAHBBY010000070.1 | Maternal | 385146 | 386977 | Green |
| HG00733 | JAHEPQ010000104.1 | Paternal | 385962 | 387799 | Red |
| HG00733 | JAHEPP010000065.1 | Maternal | 22366148 | 22367979 | Green |
| HG00735 | JAHBCH010000040.1 | Paternal | 13122143 | 13123980 | Red |
| HG00735 | JAHBCG010000054.1 | Maternal | 386159 | 387702 | Black |
| HG00741 | JAHALY010000100.1 | Paternal | 3326151 | 3327988 | Red |
| HG00741 | JAHALX010000114.1 | Maternal | 14867849 | 14869686 | Red |
| HG01071 | JAHBCF010000102.1 | Paternal | 19037145 | 19038982 | Red |
| HG01071 | JAHBCE010000099.1 | Maternal | 381021 | 382852 | Green |
| HG01106 | JAHAMC010000037.1 | Paternal | 13111824 | 13113367 | Black |
| HG01106 | JAHAMB010000077.1 | Maternal | 12036638 | 12038181 | Black |
| HG01109 | JAHEPA010000196.1 | Paternal | 384340 | 385883 | Black |
| HG01109 | JAHEOZ010000253.1 | Maternal | 383584 | 384980 | Blue |

| | | | | | |
|---|---|---|---|---|---|
| HG01123 | JAGYYZ010000097.1 | Paternal | 1592094 | 1593931 | Red |
| HG01123 | JAGYYY010000053.1 | Maternal | 385696 | 387527 | Green |
| HG01175 | JAHAMA010000163.1 | Paternal | 383013 | 384844 | Green |
| HG01175 | JAHALZ010000161.1 | Maternal | 384201 | 385744 | Black |
| HG01243 | JAHEOY010000049.1 | Paternal | 12488178 | 12489721 | Black |
| HG01243 | JAHEOX010000125.1 | Maternal | 385738 | 387569 | Green |
| HG01258 | JAGYYV010000102.1 | Paternal | 384743 | 386580 | Red |
| HG01258 | JAGYYU010000131.1 | Maternal | 383686 | 385523 | Red |
| HG01358 | JAGYZB010000111.1 | Paternal | 386131 | 387674 | Black |
| HG01358 | JAGYZA010000059.1 | Maternal | 3332334 | 3333877 | Black |
| HG01361 | JAGYYX010000045.1 | Paternal | 385392 | 387229 | Red |
| HG01361 | JAGYYW010000050.1 | Maternal | 17469033 | 17470576 | Black |
| HG01891 | JAGYVO010000053.1 | Paternal | 21524600 | 21526143 | Black |
| HG01891 | JAGYVN010000056.1 | Maternal | 24497942 | 24499773 | Green |
| HG01928 | JAGYVQ010000038.1 | Paternal | 385685 | 387228 | Black |
| HG01928 | JAGYVP010000046.1 | Maternal | 25668786 | 25670329 | Green |
| HG01952 | JAHAME010000032.1 | Paternal | 386538 | 388375 | Red |
| HG01952 | JAHAMD010000048.1 | Maternal | 386328 | 388159 | Black |
| HG01978 | JAGYVS010000006.1 | Paternal | 383134 | 384677 | Black |
| HG01978 | JAGYVR010000105.1 | Maternal | 383133 | 384676 | Black |
| HG02055 | JAHEPK010000109.1 | Paternal | 12099561 | 12101392 | Green |
| HG02055 | JAHEPJ010000128.1 | Maternal | 385702 | 387245 | Black |
| HG02080 | JAHEOW010000012.1 | Paternal | 386450 | 388281 | Green |
| HG02080 | JAHEOV010000101.1 | Maternal | 386478 | 388309 | Green |
| HG02109 | JAHEPG010000044.1 | Paternal | 385169 | 386712 | Black |
| HG02109 | JAHEPF010000205.1 | Maternal | 386657 | 388200 | Black |
| HG02145 | JAHKSG010000015.1 | Paternal | 16426311 | 16427854 | Black |
| HG02145 | JAHKSF010000110.1 | Maternal | 18842194 | 18843737 | Black |
| HG02148 | JAHAMG010000107.1 | Paternal | 386171 | 388002 | Green |
| HG02148 | JAHAMF010000035.1 | Maternal | 386204 | 388041 | Red |
| HG02257 | JAGYVI010000052.1 | Paternal | 8775388 | 8776931 | Black |
| HG02257 | JAGYVH010000008.1 | Maternal | 384860 | 386691 | Green |
| HG02486 | JAGYVM010000070.1 | Paternal | 10006631 | 10008174 | Black |
| HG02486 | JAGYVL010000054.1 | Maternal | 13134792 | 13136335 | Black |
| HG02559 | JAGYVK010000090.1 | Paternal | 384282 | 386113 | Green |
| HG02559 | JAGYVJ010000046.1 | Maternal | 384125 | 385521 | Blue |
| HG02572 | JAHAOW010000228.1 | Paternal | 2534622 | 2536165 | Black |
| HG02572 | JAHAOV010000285.1 | Maternal | 2159093 | 2160636 | Black |
| HG02622 | JAHAOO010000001.1 | Paternal | 382951 | 384494 | Black |
| HG02622 | JAHAON010000008.1 | Maternal | 13136641 | 13138184 | Black |
| HG02630 | JAHAOQ010000017.1 | Paternal | 381740 | 383283 | Black |

| | | | | | |
|---|---|---|---|---|---|
| HG02630 | JAHAOP010000028.1 | Maternal | 16114869 | 16116412 | Black |
| HG02717 | JAHAOS010000070.1 | Paternal | 8812957 | 8814212 | Black |
| HG02717 | JAHAOR010000025.1 | Maternal | 13118906 | 13120449 | Black |
| HG02723 | JAHEOU010000082.1 | Paternal | 18888872 | 18890415 | Black |
| HG02723 | JAHEOT010000245.1 | Maternal | 3328415 | 3329958 | Black |
| HG02818 | JAHEOS010000044.1 | Paternal | 383235 | 385066 | Green |
| HG02818 | JAHEOR010000191.1 | Maternal | 385402 | 386945 | Black |
| HG02886 | JAHAOU010000095.1 | Paternal | 16591016 | 16592559 | Black |
| HG02886 | JAHAOT010000063.1 | Maternal | 25052267 | 25053810 | Black |
| HG03098 | JAHEPM010000128.1 | Paternal | 13777298 | 13778841 | Black |
| HG03098 | JAHEPL010000029.1 | Maternal | 387329 | 388872 | Black |
| HG03453 | JAGYVW010000042.1 | Paternal | 3340771 | 3342314 | Black |
| HG03453 | JAGYVV010000140.1 | Maternal | 384893 | 386724 | Green |
| HG03486 | JAHEOQ010000031.1 | Paternal | 8841493 | 8843324 | Green |
| HG03486 | JAHEOP010000085.1 | Maternal | 385591 | 387134 | Black |
| HG03492 | JAHEPI010000021.1 | Paternal | 18267643 | 18269186 | Black |
| HG03492 | JAHEPH010000001.1 | Maternal | 384547 | 386378 | Green |
| HG03516 | JAGYYT010000034.1 | Paternal | 24857464 | 24859007 | Black |
| HG03516 | JAGYYS010000007.1 | Maternal | 25660968 | 25662511 | Black |
| HG03540 | JAGYVY010000133.1 | Paternal | 15842834 | 15844377 | Black |
| HG03540 | JAGYVX010000095.1 | Maternal | 15680201 | 15681744 | Black |
| HG03579 | JAGYVU010000033.1 | Paternal | 385111 | 386654 | Black |
| HG03579 | JAGYVT010000054.1 | Maternal | 384988 | 386819 | Green |
| NA18906 | JAHEOO010000012.1 | Paternal | 13131597 | 13133140 | Black |
| NA18906 | JAHEON010000061.1 | Maternal | 385316 | 387147 | Green |
| NA20129 | JAHEPE010000097.1 | Paternal | 3335060 | 3336891 | Green |
| NA20129 | JAHEPD010000165.1 | Maternal | 384671 | 386502 | Green |
| NA21309 | JAHEPC010000285.1 | Paternal | 5591544 | 5593375 | Green |
| NA21309 | JAHEPB010000234.1 | Maternal | 386377 | 388208 | Green |

**Scalability of KmerSV**

Supplementary Table 3 presents the performance metrics of KmerSV when applied to sequence pairs of varying lengths, ranging from approximately one thousand to one million base pairs. These evaluations were performed using a single-threaded configuration. KmerSV consistently exhibits efficient runtime performance, scaling linearly with the length of the genome sequences. Notably, the tool is engineered for scalability; it can partition the reference sequence into smaller segments and utilize multi-threading to accelerate the analysis process. Additionally, KmerSV allows for the extraction of 31-mers at user-defined interval lengths instead of every 31-mers, further enhancing processing speed for large genomic datasets. With its built-in scalability features, KmerSV holds the potential to analyze exceedingly long sequences, efficiently pinpointing regions that may contain structural variations.

Supplementary Table 3: Running time of KmerSV across varying sequence lengths.

| Genome Length (base pairs) | Running Time (seconds) |
|---|---|
| 1000 | 0.35 |
| 10,000 | 5.59 |
| 100,000 | 41.37 |
| 1,000,000 | 473.21 |
